# Portrait of a generalist bacterium: pathoadaptation, metabolic specialization and extreme environments shape diversity of *Staphylococcus saprophyticus*

**DOI:** 10.1101/2023.08.18.553882

**Authors:** Madison A. Youngblom, Madeline R. Imhoff, Lilia M. Smyth, Mohamed A. Mohamed, Caitlin S. Pepperell

## Abstract

*Staphylococcus saprophyticus* is a Gram-positive, coagulase-negative staphylococcus found in diverse environments including soil and freshwater, meat, and dairy foods. *S. saprophyticus* is also an important cause of urinary tract infections (UTIs) in humans, and mastitis in cattle. However, the genetic determinants of virulence have not yet been identified, and it remains unclear whether there are distinct sub-populations adapted to human and animal hosts. Using a diverse sample of *S. saprophyticus* isolates from food, animals, environmental sources, and human infections, we characterized the population structure and diversity of global populations of *S. saprophyticus*. We found that divergence of the two major clades of *S. saprophyticus* is likely facilitated by barriers to horizontal gene transfer (HGT) and differences in metabolism. Using genome-wide association study (GWAS) tools we identified the first Type VII secretion system (T7SS) described in *S. saprophyticus* and its association with bovine mastitis. Finally, we found that in general, strains of *S. saprophyticus* from different niches are genetically similar with the exception of built environments, which function as a ‘sink’ for *S. saprophyticus* populations. This work increases our understanding of the ecology of *S. saprophyticus* and of the genomics of bacterial generalists.

**Data summary:** Raw sequencing data for newly sequenced *S. saprophyticus* isolates have been deposited to the NCBI SRA under the project accession PRJNA928770. A list of all genomes used in this work and their associated metadata are available in the supplementary material. Custom scripts used in the comparative genomics and GWAS analyses are available here: https://github.com/myoungblom/sapro_genomics.

**Impact statement:** It is not known whether human and cattle diseases caused by *S. saprophyticus* represent spillover events from a generalist adapted to survive in a range of environments, or whether the capacity to cause disease represents a specific adaptation. Seasonal cycles of *S. saprophyticus* UTIs and molecular epidemiological evidence suggest that these infections may be environmentally-acquired rather than via transmission from person to person. Using comparative genomics and genome wide association study tools, we found that *S. saprophyticus* appears adapted to inhabit a wide range of environments (generalist), with isolates from animals, food, natural environments and human infections being closely related. Bacteria that routinely switch environments, particularly between humans and animals, are of particular concern when it comes to the spread of antibiotic resistance from farm environments into human populations. This work provides a framework for comparative genomic analyses of bacterial generalists and furthers our understanding of how bacterial populations move between humans, animals, and the environment.

## Introduction

*Staphylococcus saprophyticus* is a Gram-positive, coagulase-negative staphylococcus (CNS) that is a major cause of urinary tract infections (UTIs) in reproductive aged women (*1*). *S. saprophyticus* is also found in a wide variety of other niches including natural environments like soil, fresh and salt water (*2*, *3*). *S. saprophyticus* colonizes and infects animals (*4*, *5*) where it can cause bovine mastitis (*6*), and is found in food processing environments and animal food products (*7–10*). Despite its relevance to human and animal health, little is known about the factors – host and bacterial – that influence *S. saprophyticus* infections.

Previous genomic studies have shown that bacterial isolates from different sources are intermingled on the phylogeny (*10–12*). Prior to the widespread use of whole-genome sequencing (WGS), pulsed field gel-electrophoresis (PFGE) genotyping of *S. saprophyticus* isolates causing UTIs showed that diverse strain types can cause infection in humans (*13*, *14*). Genomic surveys of putative virulence factors in *S. saprophyticus* from different sources show similar distributions of putative virulence genes, particularly adhesins that enable colonization of the human urinary tract (*15*, *16*). A recent genomics study showed that *S. saprophyticus* from meat processing plants have high genetic relatedness to human UTI isolates from surrounding communities (*10*). These results demonstrate that diverse *S. saprophyticus* strains cause disease in humans, and prior studies have failed to identify virulence factors or transmission barriers that separate pathogenic from non-pathogenic strains. Thus, it is not clear whether sub-populations of *S. saprophyticus* are specifically adapted to cause disease, or more generally, if sub-populations of *S. saprophyticus* are uniquely adapted to the various niches they inhabit.

In order to address this knowledge gap, we performed genome wide association studies (GWAS) of the largest and most diverse sample of *S. saprophyticus* genomes analyzed to date. We used comparative genomics to characterize the diversity and population structure of *S. saprophyticus* and revealed genetic and ecological factors driving divergence between the two major clades of *S. saprophyticus*. GWAS investigations of genomic signatures of host and niche adaptation show that the majority of *S. saprophyticus* isolates appear to be generalists moving freely between environments. We identified exceptions to genomic generalism among bacteria inhabiting built environments and those causing bovine mastitis.

## Results

Combining newly sequenced isolates as well as all data publicly available at the time of analysis, we produced a sample of 780 *Staphylococcus saprophyticus* genomes. Genomes were categorized by isolation source into the following groups (see Methods): animal, built environment, food, human, and natural environment (Supplementary Data 1).

### Diversity among S. saprophyticus populations

A phylogeny inferred from an alignment of core genome sequences shows that isolates from diverse sources are closely related, with no evidence of a particular sub-population or phylogenetic lineage being adapted to a single niche (Figure 1). This contrasts with other bacterial species with multiple niches where adaptation to a particular host or environment is more obvious and specific lineages are associated with specific hosts, as is the case for *Staphylococcus aureus* (*17*). The lack of geographic and temporal structure on the phylogeny was striking, with examples of bacteria from different continents and/or isolated decades apart being very closely related (Figure S1). For example, one strain from the Washington state Pacific Ocean isolated in 2008 is separated by only 47 core genome SNPs (5e-5 SNPs per site) from a strain isolated from a Norwegian bathroom in 2021 (Figure S1).

**Figure 1:**
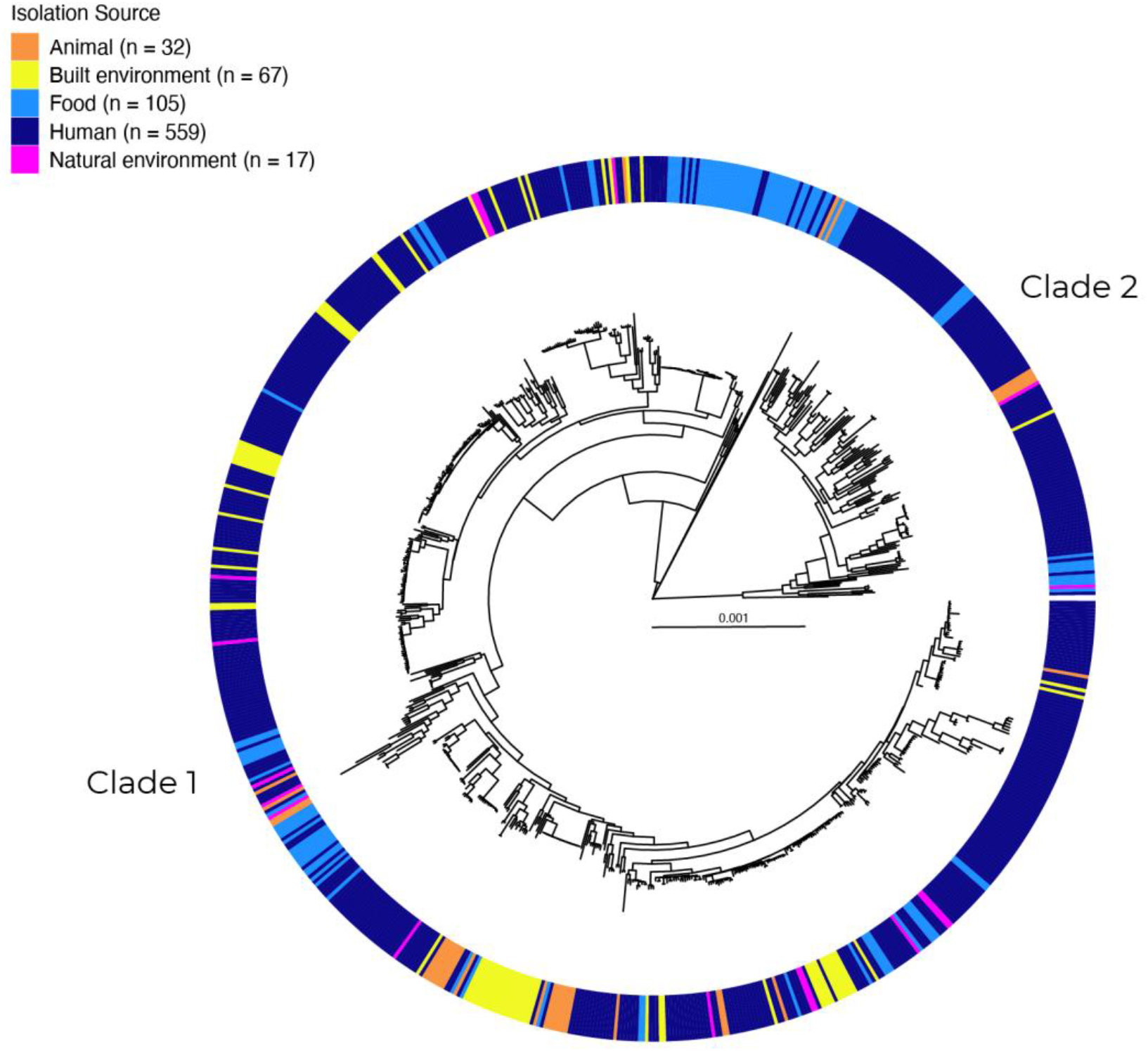
*S. saprophyticus* from diverse niches are intermingled on the phylogeny. A phylogeny was inferred from an alignment of core genes using RAxML and recombinant regions removed with ClonalFrameML. Isolates form two distinct clades: Clade 1 (n = 646) is more common in our sample with ∼5x more isolates than Clade 2 (n = 134). Isolation sources indicated on outer ring. Scale bar represents nucleotide substitutions per site.

Among an average of 2,461 genes per genome, 85% of genes in each isolate are core genes, with 15% categorized as accessory genes (Table S1). We found that despite the high proportion of core genes in each genome, isolates in our sample showed extremely high diversity in accessory gene content: the pangenome is made up of ∼14,000 genes, 80% of which are present in less than 15% of isolates in our sample (Table S1). This indicates that although a high proportion of the *S. saprophyticus* genome is conserved within the species, the remaining gene content is made up of largely unique genes, likely acquired from other species via horizontal gene transfer (HGT).

To investigate the role of HGT in shaping *S. saprophyticus* populations we used ClonalFrameML (*18*) to infer recombinant fragments within the core genome and calculate the relative contributions of recombination and mutation to observed genetic diversity (r/m). We found that *S. saprophyticus* has an r/m value of 1.2, similar to *S. aureus*, which was recently reported to have a core-genome r/m of ∼1 (*19*). Compared to other species with wide host-ranges such as *Campylobacter jejuni* (r/m = 150) and *Listeria monocytogenes* (r/m = 85) (*19*), it appears that HGT has a less prominent role in diversification of the *S. saprophyticus* core genome. Plasmids are an alternative mechanism for introducing genetic novelty, via acquisition of accessory gene content. We used a comprehensive database of plasmid sequences (*20*) to identify plasmids in our de novo assemblies and found that plasmid content was variable among isolates of *S. saprophyticus.* About half the isolates in our study did not have any matches to the plasmid database, while the other half of isolates had between one and five plasmids (Figure S2). We used multi-dimensional scaling (MDS) to group plasmid sequences into eight different sequence groups and found these groups distributed throughout the phylogeny (Figure S2). Overall, this indicates that like the patterns we observed in accessory gene content, plasmid content is highly variable between strains and likely represents an important source of genetic novelty for *S. saprophyticus* given the relatively low rates of core genome HGT. Given the diversity in plasmid content we observed here it is possible that isolates without any identifiable plasmids carry plasmids that do not share sequence similarity with those in the database. Future work using long-read sequencing will elucidate the true diversity in *S. saprophyticus* plasmid content.

### Barriers to horizontal gene transfer between major clades of S. saprophyticus

The presence of two clades within the global population of *S. saprophyticus*, designated here as Clades 1 and 2, has been described in previous genomic surveys (*10–12*). These clades are genetically distinct at the core genome level – between-clade ANI values range from 95-99% - but not diverged enough to be considered separate sub-species by the standard definition (Figure S3A). Although strains from the two clades appear to be found from similar environments (Figure 1), similar geographic regions, and similar time periods (Figure S1), our results demonstrate that the clades occupy distinct, cryptic niches.

We performed separate pangenome analyses on each clade to identify whether the clades were uniform with respect to accessory gene content and found that 1) the clades have different pangenome structures and 2) the clades contain distinct accessory gene content. Rarefaction and accumulation curves show that, adjusting for differences in sample size, Clade 1 has a larger pangenome despite similarly sized core genomes (Figure 2A). We also examined the frequencies of accessory genes and found that they are maintained at different frequencies in the two clades, consistent with barrier(s) to gene flow between the clades (Figure 2B). When examining genes that are shared across clades, including both core and accessory genes, we found nucleotide diversity to be significantly higher for between-clade versus within-clade comparisons (Figure S3B). This provides further evidence of barriers to gene flow between clades. The two clades are distinct with respect to pan genome size, nucleotide sequence of core and accessory genes, and content of the accessory genome.

It was previously reported that patterns of recombination differed between the clades (*10*), a pattern that was replicated in our study: Clade 2 has a 3x higher r/m value as well as significantly more and significantly longer recombinant fragments than Clade 1 (Figure S4). Despite having a higher recombination rate, Clade 2 has a smaller pangenome than Clade 1 (Figure 2A). Differences in pangenome structure and recombination within the same species can indicate differences in bacterial niche (*21*, *22*). We used FastGear (*23*) to detect HGT between the two clades, and these analyses identified very few inter-clade recombination events (Figure 2C). Taken together, these analyses suggest that the clades occupy subtly different niches and that one or more barriers to HGT prevent genetic exchange between the clades.

Restriction-modification systems (RMS) serve as innate bacterial immune systems by eliminating foreign DNA based on methylation patterns (*24*) and are a mechanism of preventing DNA exchange. We used a database of RM system genes (*25*) to annotate RMS genes in our sample and identified a set of RM genes at substantially different frequencies in each clade (Figure S5). These differences in RMS may provide a mechanistic barrier to HGT between clades, however are likely to be other factors that maintain differences in RMS between the clades. We hypothesized that differences in metabolism between isolates from different clades would allow them to co-localize yet occupy distinct ecological niches. We used the program Metabolic (*26*) to annotate metabolic pathways and enzymes in our assemblies, and found that the gene encoding beta-galactosidase *(ebgA,* similar to gene SSP0105 in the reference genome), an enzyme involved in lactose metabolism, is differentially maintained within clades of *S. saprophyticus* with 97% of Clade 1 isolates carrying the gene, compared to only 30% of Clade 2 isolates (Figure 3). We used growth on differential medium (MacConkey agar) to test the capacity for lactose metabolism in isolates from the two clades. All of the Clade 1 isolates in our strain collection encode *ebgA*, and all isolates that we tested from this clade were able to metabolize lactose (Figure 3). For Clade 2, we were able to test strains with and without beta-galactosidase (beta-gal^+/-^). As predicted, beta-gal^−^ strains were defective for lactose metabolism. One of two beta-gal^+^ Clade 2 isolates was able to metabolize lactose. The Clade 2 strain C085 (human UTI) encodes a full-length copy of *ebgA*, without any defects in sequence or length, yet did not metabolize lactose (Figure 3). In summary, our data reveal differences in metabolism between the two major clades of *S. saprophyticus*, with Clade 2 isolates commonly lacking the capacity for lactose metabolism through the absence *of ebgA* and other mechanisms. We hypothesize that the genetic barrier we identified between clades reflects mechanistic barriers to between-clade HGT via differentiated RMS and their separation into distinct metabolic niches.

**Figure 2:**
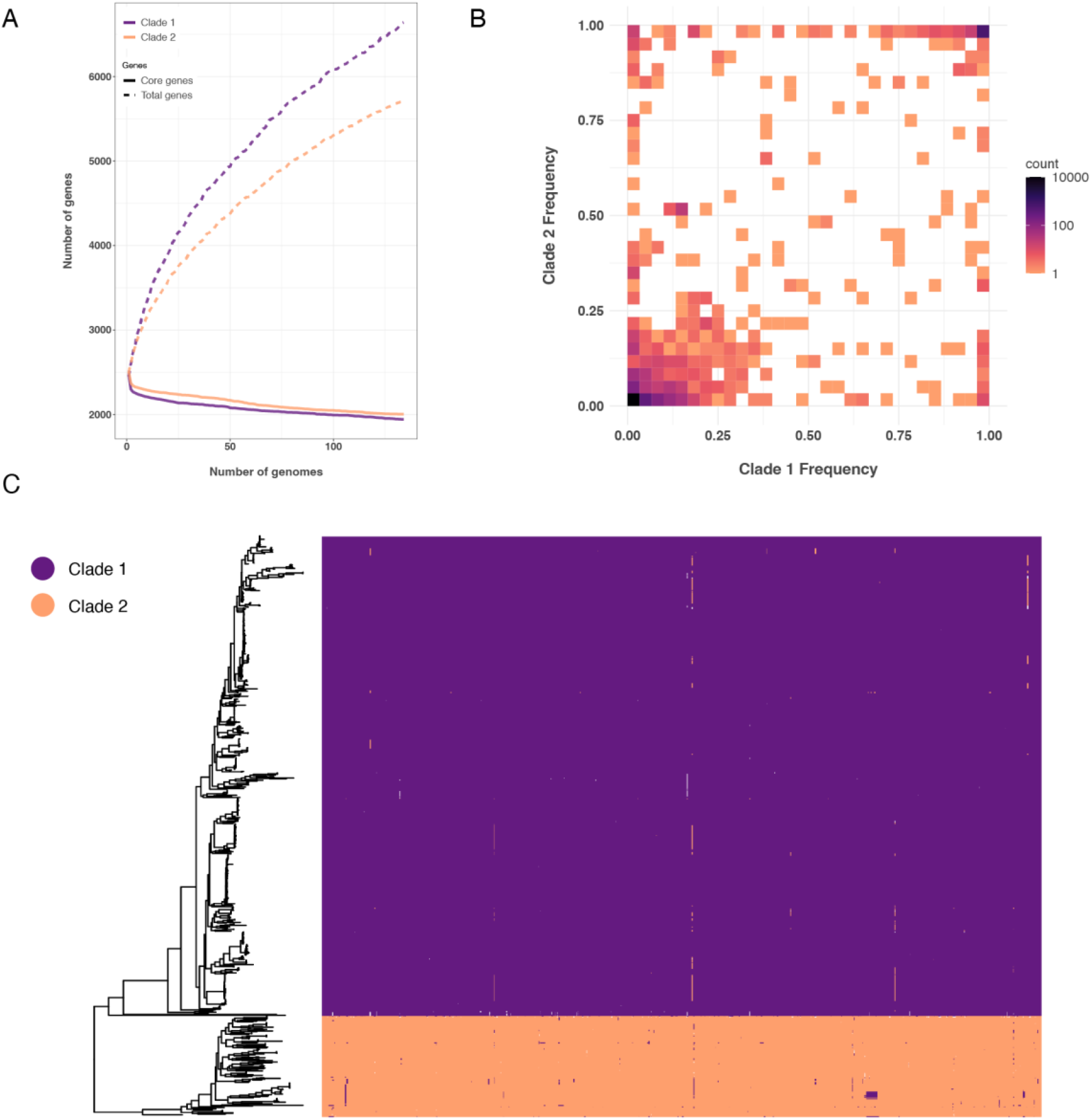
*S. saprophyticus* clades are reproductively isolated. A) Accumulation and rarefaction curves calculated using gene presence/absence matrices from separate pangenome analyses of each clade. Clade 1 has a larger accessory genome than Clade 2, while core genome sizes of the two clades are similar. B) Accessory genes present in at least one isolate of each clade are plotted to compare their frequency, showing that accessory gene content is differentially maintained by the two clades. C) Inter-clade recombination predicted using FastGEAR. Despite evidence of moderate intra-clade recombination, inter-clade recombination is rare, indicating that the clades are reproductively isolated.

### Pathoadaptation of S. saprophyticus associated with bovine mastitis

While the genetic differentiation of the two major clades appears to reflect an important separation of metabolic niches, the clade structure does not explain bacterial associations with pathogenicity or other traits as Clades 1 and 2 are both found in diverse environments, including within humans and other animals (Figure 1). In order to identify genotype-phenotype associations with the varied environments where *S. saprophyticus* is found, we first examined accessory gene content. Broad-scale patterns of accessory gene presence/absence are for the most part homogenous across isolation sources (Figure 4A). Overall differences in accessory gene frequency are minimal and appear random, except for a handful of genes that were uniquely associated with animal isolates. Using Scoary (*27*) to test the strength of association between accessory gene content and isolation source we found that these genes were highly significantly associated with animal isolates (Supplementary Data 2). Closer inspection of the genes revealed a full type VII secretion system (T7SS) operon (Figure 4B). This is the first description of a T7SS in *S. saprophyticus*, which has been described previously in other coagulase-negative staphylococci (CNS) (*28*, *29*). In our sample of *S. saprophyticus* the T7SS is almost exclusively found in isolates from bovine mastitis: 78% (14/18) bovine mastitis isolates in our sample carry the T7SS, while only one non-mastitis strain (19-02, human UTI) carries the element (Figure 4B). The *S. saprophyticus* T7SS genes are organized in an operon structure very similar to that of *S. aureus* (*30*) and *S. lugdunensis* (*28*) that is conserved across isolates in this sample. Blast results show that the sequence of the T7SS in our sample most closely resembles T7SS genes from *S. arlettae*, a closely related CNS species that colonizes animals (*31*). Given its distribution across the phylogeny, we hypothesize that the T7SS has been horizontally acquired multiple times from other *Staphylococcus* spp. and that it is under positive selection in *S. saprophyticus*. Given the association with mastitis isolates, we further hypothesize that it plays a role in mastitis pathogenesis; virulence properties conferred by the T7SS could be advantageous or they may represent off target effects. In *S. aureus* the T7SS is required for virulence in many models of infection (*32*) and is important for resistance against host-immune pressures (*33*). The same may be true for *S. saprophyticus*, perhaps allowing bacteria to invade otherwise depauperate bovine tissues and escape from competition with other microbes. The T7SS provides the clearest example in our data of a significant association between accessory gene content and host or environmental niche, which may be a function of incomplete sampling and/or a multitude of adaptive pathways for *S. saprophyticus* to the same niche.

**Figure 3:**
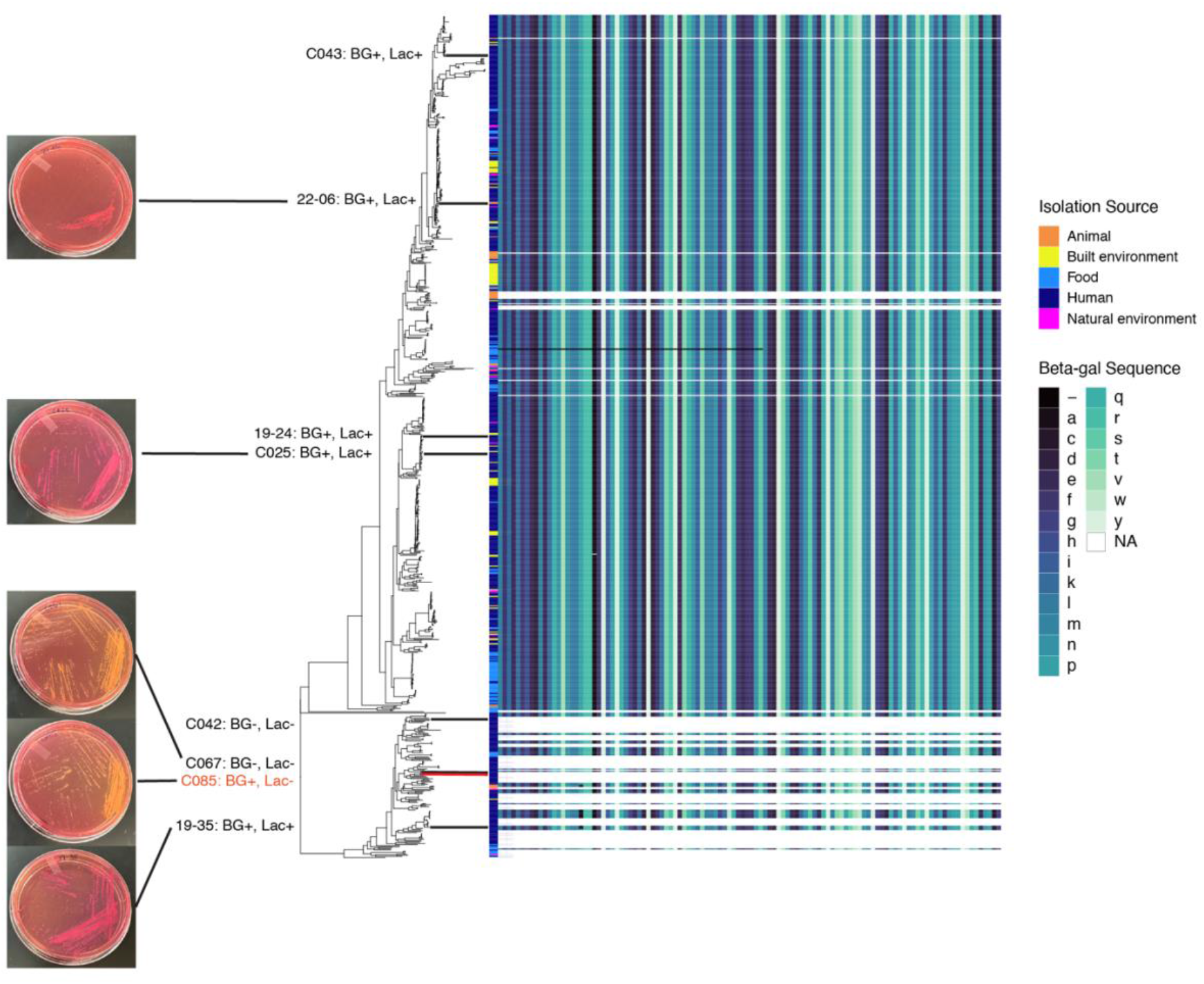
Niche separation of *S. saprophyticus* clades by lactose metabolism. Amino acid sequence of EbgA beta-galactosidase (beta-gal) is plotted next to the core genome phylogeny showing that the sequence is highly conserved at the protein level. Beta-gal is almost fixed in Clade 1 isolates (97%), while a minority of Clade 2 isolates carry the gene (30%). Lines to the right of the phylogeny indicate the strains that were tested for their ability to metabolize lactose. BG+/- indicates the presence/absence of beta-gal in that strain, and Lac+/- indicates whether that strain was positive/negative for lactose metabolism on MacConkey agar. Representative photos from each clade show results positive for lactose metabolism (Lac+, pink colonies) and those negative for lactose metabolism (Lac-, yellow colonies). Out of 8 strains we tested, only one (C085, shown in red) was an outlier in that it carries beta-gal but did not test positive for lactose metabolism.

**Figure 4:**
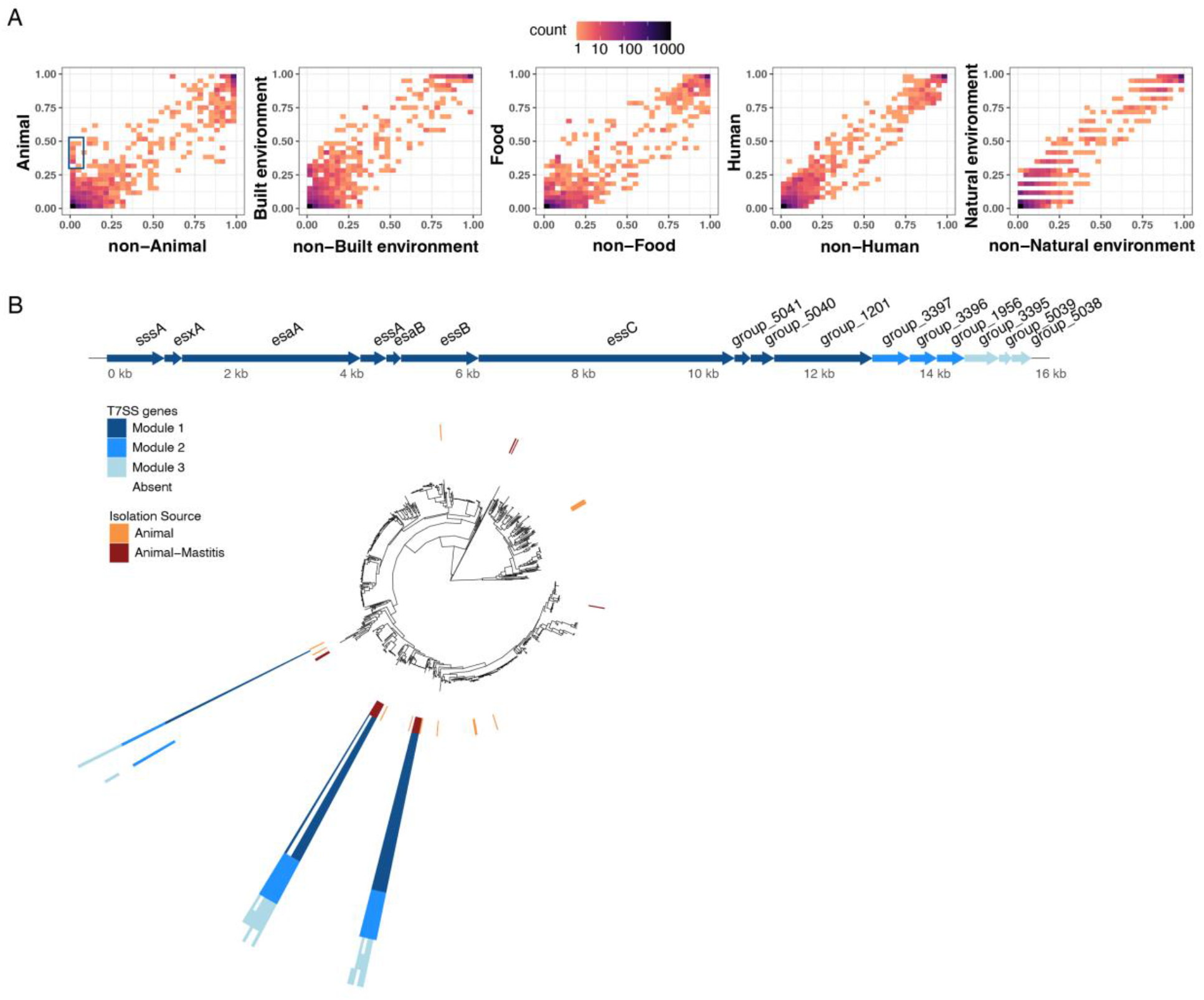
Type VII secretion system found in bovine mastitis isolates. A) Frequencies for all accessory genes in the pangenome (n = 11,952) are plotted by isolation source on the y-axis with the average frequency in all other sources on the x-axis. Overall, accessory genes are at similar frequencies in all niches, except for a few genes that appeared uniquely associated with animal isolates (outlined in blue). B) Genes encoding a Type VII secretion system (T7SS) are very significantly associated with isolates from bovine mastitis. 78% (14/18) of bovine mastitis isolates encode a full or partial T7SS, while only 1 other isolate (19-02, human UTI) encodes the T7SS. Genes in the T7SS operon is divided into three “modules” that are generally found together within the same genome. Inner ring around the core genome phylogeny indicates isolates from cases of bovine mastitis as well as isolates from other animal sources. Outer rings indicate presence/absence of T7SS genes (in the same order as the gene operon) and are colored by module.

### Associations between SNPs and isolate source

To further investigate associations between bacterial genetic loci and environmental niche, we performed genome-wide association studies (GWAS) of single nucleotide polymorphisms (SNPs). We used pySEER (*34*), a tool specifically designed for use with microbial genomes, to test whether specific genome variants were associated with adaptation to a particular niche. Two highly significant results include a synonymous SNP associated with natural environments that lies within an L-serine dehydratase, and another synonymous SNP associated with animals that lies within the 2-component system regulator YycH, which in *S. aureus* helps regulate the expression of autolysins (*35*). Aside from these two examples, out of ∼6,500 variants we tested from the whole genome alignment, generally few if any variants were significantly associated with a single isolation source (Figure 5). This pattern also held true when the same analysis was performed using only core genome SNPs (Figure S6). Built environments were, however, exceptional among isolate sources. A markedly larger number of variants associated with isolates from built environments in both whole-and core genome analyses. SNPs significantly associated with built environments make up 75% and 80% of all significant SNPs from the whole-genome and core genome analyses, respectively. Examining the types of mutations associated with isolates from built environments, we find that a large proportion of them are within coding regions, either synonymous or non-synonymous, indicating that many of these mutations could be expected to have impacts on protein function (Figure 5B). We annotated the genes containing these associated variants using clusters of orthologous group (COG) categories and found that most are either function unknown (S) or amino acid metabolism (E) (Table S2).

**Figure 5:**
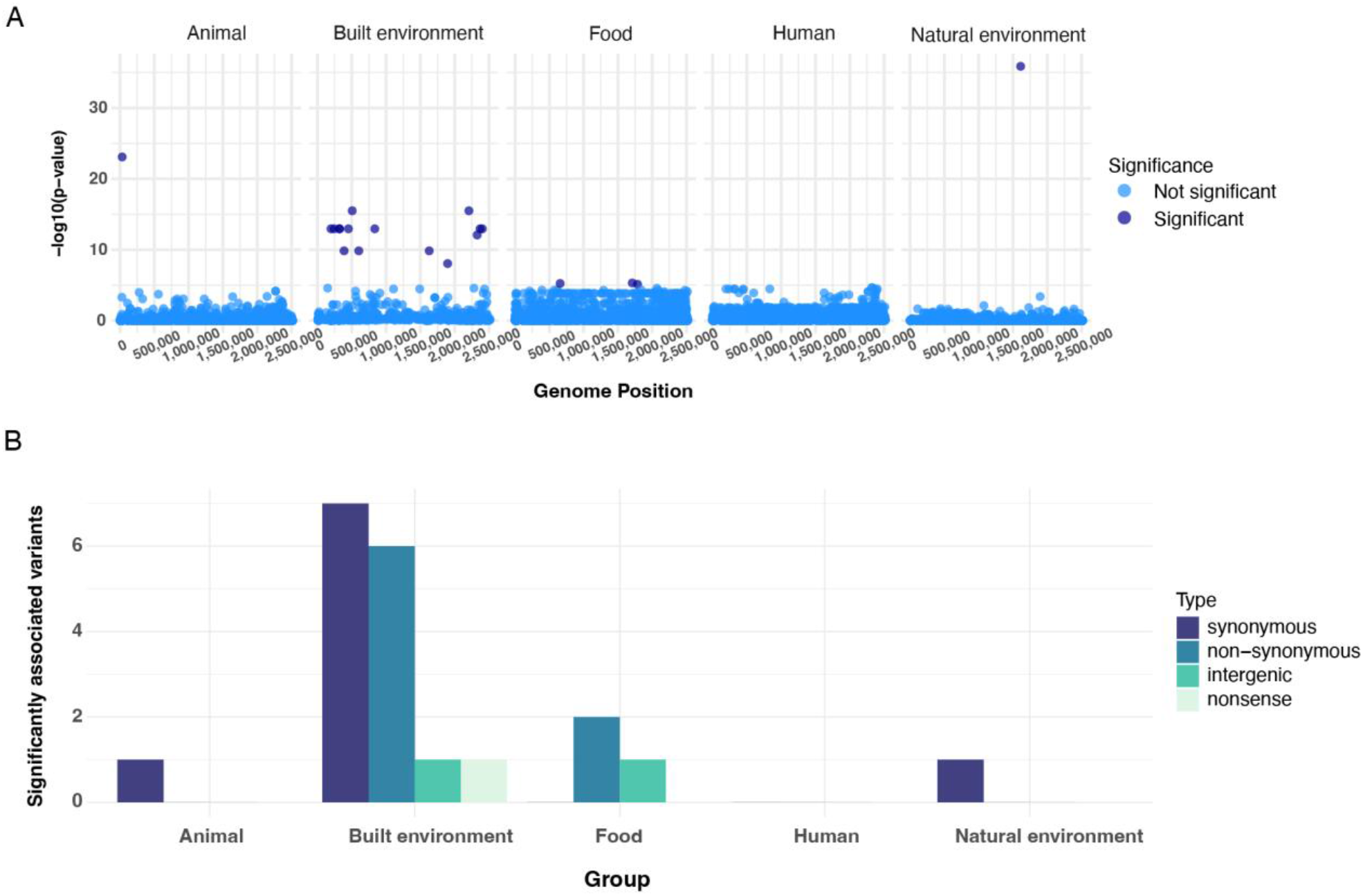
Few whole-genome variants associated with niche adaptation. A) GWAS on all bi-allelic SNPs from whole genome alignment (n = 6,457) was performed using pySEER with a linear mixed model. SNPs identified as significantly associated with a particular niche are in dark blue, non-significant in light blue. Isolates from built environments have more variants that distinguish them from other isolates: 75% of all significant SNPs were associated with built environments. B) Number of significant pySEER SNPs plotted by associated source and mutation type. SNPs associated with isolates from built environments are relatively skewed towards non-synonymous variants.

### Built environments exert unique selective pressure

We hypothesized that the striking number of *S. saprophyticus* SNPs associated with built environments pattern could result from unique selective pressures encountered in this niche. We first examined overall ratios of non-synonymous (dN/dS) and intergenic (dI/dS) variation to synonymous variation within sub-populations from each source and found that, apart from natural environments, different source types had similar values of genome-wide dN/dS. Natural environments had significantly higher dN/dS and dI/dS (Figure 6A). We hypothesize that this pattern could reflect the increased relative diversity of environments we have collected under “natural environments” (including soil, salt and fresh water) but future analyses of a larger sample of isolates from natural environments may reveal more.

**Figure 6:**
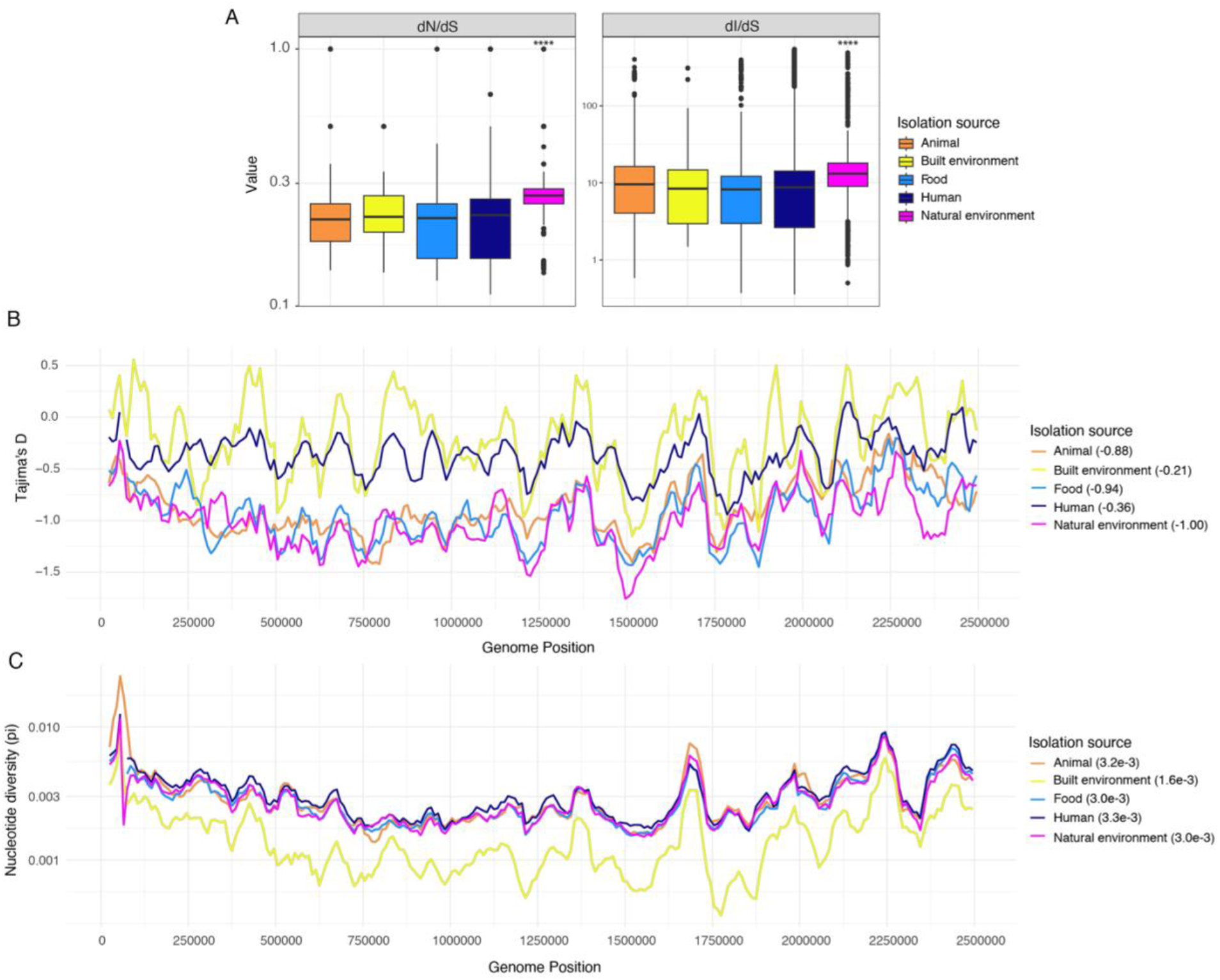
Niches of *S. saprophyticus* exert different selective pressures. A) dN/dS (left) and dI/dS (right) were calculated for all pairs of core genome alignments from a given isolation source. Isolates from natural environments have significantly (Mann-Whitney U test with Bonferroni correction, ****: *P* < 0.0001) higher dN/dS and dI/dS than other niches. Population genetics statistics were calculated in sliding-windows (window size: 50,000 bp, step size: 10,000 bp) across whole genome alignments of isolates from different sources. Alignments were repeatedly (100x) sub-sampled to the size of the smallest sample (natural environment, n=17) and the mean Tajima’s D (C) and nucleotide diversity (D) values from all sub-samples per window was plotted. Mean genome-wide statistics are listed in the legend of each plot.

An alternative, and not mutually exclusive, explanation could lie in the complexity of natural environments such as soil, which is estimated to contain a majority of the earth’s biodiversity (*36*). Built environments stood out from the other sources in our sample with respect to other aspects of population genetic diversity summarized with Tajima’s D (the difference between the observed and expected variation within a population) and nucleotide diversity (pi). The bacterial sub-population from built environments has a neutral Tajima’s D (D = 0), consistent with a neutrally evolving population with stable population size (Figure 6C) and lower nucleotide diversity, indicating a more genetically homogeneous population (Figure 6D). This contrasts with isolates from animals, food and natural environments, which have higher diversity and negative genome-wide values of Tajima’s D, consistent with population expansion. In summary, we found that isolates from built environments have signatures of a stable population size, low diversity, and more significantly associated variants that distinguish them from other sources. We hypothesize that these results indicate a non-random filtering of isolates that can survive in built environments, leading to a relatively stable population size and reduced diversity.

## Discussion

Analyzing a diverse sample of genomes, we have identified barriers to horizontal gene transfer (HGT) and differences in metabolic capacity between the major clades of *S. saprophyticus*. Although the division into two clades is fundamental to the genetic structure of *S. saprophyticus* populations, it does not explain niche associations for this peripatetic bacterium. *S. saprophyticus* appears to be panmictic: diverse bacteria are associated with individual environments, and conversely, diverse environments are associated with genetically similar bacteria. Within this fluid population structure, our fine-scale analyses have revealed genomic imprints of specific environments, namely pathoadaptation via the acquisition of a Type VII secretion system associated with bovine mastitis, and an overall winnowing of diversity in association with what are likely extreme environments, i.e. human-made environments. Overall, this work paints a picture of a bacterium that is both generally adapted to transition between diverse environments and also adapt to specific niches.

### Divergence of S. saprophyticus clades

Our results show that *S. saprophyticus* isolates from Clades 1 and 2 are genetically distinct with respect to gene content (Figure 2B) and nucleotide sequence of the core and accessory genomes (Figure S3). We further find evidence suggesting they are reproductively isolated, as recombination between clades appears to be rare (Figure 2C). Differences in restriction-modification systems (RMSs) (Figure S5) and metabolic capacity (Figure 3) offer potential explanations for the observed clade structure, which is likely due to multiple mechanistic and ecological factors, including the reinforcing effect of genetic differentiation in depressing recombination (*37*). These results parallel observations in other species of coagulase negative *Staphylococcus*: distinct clades of *Staphylococcus epidermidis* that appear specialized to different niches on the human body show evidence of barriers to HGT in their genomes (*38*). However, unlike *S. epidermidis*, our data suggest that *S. saprophyticus* isolates from each clade inhabit the same environments (Figure 1), excluding the possibility of a simple spatial barrier to HGT between clades. Other examples of barriers to HGT between isolates of the same species include *Campylobacter jejuni* (*39*) and species within the genus *Gardnerella* (*40*) which are both highly recombinogenic and yet display significant restrictions on HGT in natural populations. In neither case was a mechanistic barrier to HGT identified; in fact, *C. jejuni* lineages are able to exchange genetic material in vitro, indicating that the barriers to HGT in natural populations are ecological.

### Diverse ecology and generalism of S. saprophyticus

*S. saprophyticus* is not a permanent member of the human genitourinary or gastrointestinal tracts but seems to be a transient colonizer in a minority of the population (*41*). A striking feature of colonization and infection with *S. saprophyticus* is the pattern of seasonality found in many parts of the world. In temperate climates, colonization, and infection by *S. saprophyticus* is most common in the warmer months of the year, spring through autumn (*41–44*). This distinguishes *S. saprophyticus* from other human pathogens such as uropathogenic *E. coli* and *Staphylococcus aureus*, which have a more stable residency in the microbiome punctuated by invasion and disease (*45*, *46*). *S. saprophyticus* exhibits further singularity in that, unlike other pathogens with a variety of niches and/or multiple hosts, it does not display lineage level adaptation to specific hosts (Figure 1). This contrasts with patterns among other *Staphylococcus* spp. (*17*, *47*) and *Campylobacter* spp. (*48–50*), for which specific lineages exhibit strong associations with particular hosts. We hypothesized that adaptation to different niches may occur at a finer scale in *S. saprophyticus*, but found that in general, accessory gene content and core genome variants appear homogenous across isolates from different niches (Figure 4, Figure 5). These results suggest that *S. saprophyticus* is a generalist in both habitat and host-types. Central to the idea of bacterial niche specialization is that adaptations which provide a fitness benefit in one environment will reduce fitness in another (*51*). This does not appear to be the case for *S. saprophyticus*, which is readily isolated from multiple hosts, natural and built environments as well as food products, with no evidence of restriction to a single niche.

Within this fluid structure, we did identify a striking example of pathoadaptation in the acquisition of a type VII secretion system (T7SS) that is strongly associated with bovine mastitis (Figure 4). The T7SS has been identified in other CNS species (*28*, *29*) and has a well-characterized role in the pathogenesis of *S. aureus* (*32*, *33*), but has not been previously been identified or characterized in *S. saprophyticus.* In our analyses, the T7SS was associated with a specific pathogenic niche distinct from animals in general, from commensal isolates of cattle, and even from isolates causing other kinds of invasive disease in cattle that were also present in this sample (Supplementary Data 1). We hypothesize that mastitis infections are distinct from other invasive infections in cattle due to the formation of a contained infection within abscesses. T7SS are known to be important for abscess formation in *S. aureus* (*52*– *54*). This contrasts with uncomplicated UTIs, which are also associated with *S. saprophyticus* (*1*), but which are not closed-space infections. All of the bovine mastitis isolates carrying the T7SS in our sample were from cases of sub-clinical mastitis, indicating a less severe form of mastitis where there is a lack of visible indicators of infection (*55*). Coagulase-negative staphylococci (CNS) are major causes of both clinical and sub-clinical mastitis (*56–59*), although this varies by region and it appears that in some regions CNS are less prevalent in clinical mastitis samples (*60*). Future work may reveal the role of the T7SS in *S. saprophyticus* bovine mastitis and further clarify the association between the T7SS and sub-clinical mastitis.

### Source-sink dynamics in the ecology of S. saprophyticus

We show here that broad-scale patterns of genetic diversity in *S. saprophyticus* are relatively homogenous across distinct environments. Within this large pan-genome the high diversity of accessory gene content held at rare frequencies raises the possibility that adaptation to any one niche can proceed by a multitude of pathways, which would render identification of genotype-phenotype associations challenging (*61*). This, we hypothesize, is the reason that niche does not appear to have a prominent role in structuring accessory gene content (Figure 4). In our analysis of genome variants, we found that only isolates from built environments had more than a few significantly associated SNPs (Figure 5, Figure S6). In comparison with other niches, bacteria from built environments also have lower relative diversity and a more balanced site frequency spectrum suggesting stable as opposed to expanding population size (Figure 6). Synthesizing these observations we infer that *S. saprophyticus* entering built environments undergo a non-random filtering process. We hypothesize that this process filters for variants that increase bacterial tolerance for dry environments and desiccation, which would be a fitness advantage in the built environments sampled here (generally fomites or air; Supplementary Data 1). A helpful framework for thinking about bacterial adaptation to new habitats is the source-sink model (*62*), where the “source” population consists of the permanent niche or reservoir of a bacterial species, and the “sink” population exists in a different niche or environment and is fed by the source population. Here we are using the definition of source-sink specifically adapted to bacterial pathogens (*62*), where the establishment of a sink population is not necessarily a neutral process as is often described in classical population ecology (*63*). A relevant example of these dynamics is repeated adaptation within the FimH adhesin of uropathogenic *E. coli,* which has been hypothesized to underlie the repeated emergence of *E. coli* lineages into the urinary tract (*64*). We propose that built environments represent a sink, which is fed by one or more source populations of *S. saprophyticus*. Staphylococci are some of the most abundant members of the built microbiome, and transmission from fomites of *S. aureus* is a known pathway for strains causing human infections (*65*). Conversely, studies show that the majority of the built environment microbiome is made up of human-associated microbes (*65*), indicating that transmission of *S. saprophyticus* to built environments is most likely from human sources. A clear example of this phenomenon is the *S. saprophyticus* sample from the International Space Station (Supplementary Data 1) where transmission from the natural environment, animals and or animal food products would be virtually impossible. Other built environments such as kitchens may be colonized by *S. saprophyticus* after contact with animal food products. A more thorough sampling of different built environments may identify the sources of *S. saprophyticus* and provide insight into the frequency of transmission from the built environment back into humans.

### Adaptation of Aas

In a previous study we identified a non-synonymous SNP in the bifunctional adhesin-autolysin gene *aas* that had putatively undergone a selective sweep (*12*). The derived allele of the SNP was also significantly associated with isolates from human UTIs. In the current study, the association between the *aas* allele and UTI isolates is no longer significant. However, the evidence for a selective sweep at this locus is retained in these data; using a sample 13x larger than was used for our first analysis, we have recapitulated the dip in Tajima’s D surrounding this locus that indicates a selective sweep (Figure S7). The evidence points to a sweep in Clade 1 only, with the dip in Tajima’s D being present only in the alignment of Clade 1 isolates (Figure S7). Mapping the alleles of the *aas* locus onto the core genome phylogeny of our sample shows that the alleles are highly structured on the phylogeny: 82% of Clade 1 isolates have the derived allele while only 28% of Clade 2 isolates have it (Figure S7). While this SNP in *aas* is not associated with a particular isolation source, it appears to be under directional selection, indicating that it may play a role in the evolution of *S. saprophyticus* populations beyond any role it plays in invasion of the human urinary tract. Adaptation of Aas could be a contributing factor in what appears to be recent population expansion in Clade 1, which has overall lower Tajima’s D values and shorter branch lengths relative to Clade 2 (Figure S7). It is yet unclear how this fits in with the differences in lactose metabolism we identified between the clades but it’s possible that changes to metabolic capacity and adhesin properties allow the bacterium to migrate between environments more easily, allowing for relative population expansion. Future work looking at the evolution of Aas, possibly using a source-sink framework as was described above for the FimH adhesin in *E. coli*, could reveal more about the contribution of Aas to the evolution of *S. saprophyticus* populations.

### The problem of transmission

Our results indicate that *S. saprophyticus* isolates are able to move freely between environments. What remains unclear is exactly how *S. saprophyticus* is transmitted, and which transmission pathways lead to human and animal infections. Prior epidemiological studies have shown that swimming and occupations related to food production increase the risk of *S. saprophyticus* UTI (*66*). Genitourinary colonization by *S. saprophyticus* is transient (*41*), providing further evidence that at least some infections are environmentally acquired (rather than transmitted person-to-person). An inexplicable lack of temporal signal (Figure S1B) in the phylogeny of *S. saprophyticus* makes it difficult to ascertain the directionality of transmission between different hosts and environments using traditional phylogenetic approaches (*10–12*). Directions for new research may be taken from the study of other generalist bacterial pathogens with multiple environmental reservoirs. For example, species of the genus *Campylobacter* similarly occupy a wide variety of environments and are a leading cause of food-borne illness (*67*). Studies combining epidemiological and sequencing data have proven very useful in identifying the animal and environmental sources of *Campylobacter* infection (*68–70*). Additionally, computational models of transmission dynamics in *Campylobacter* have highlighted the role of insect vectors in transmission (*71*, *72*). For *S. saprophyticus* there is still much to learn about transmission dynamics, including the role of food products and built environments in transmitting *S. saprophyticus*. We know that occupations in food production are a risk factor for *S. saprophyticus* UTI (*66*), but this group represents a very small proportion of the population. This opens the question of what, if any role, routine contact with and consumption of animal food products plays in *S. saprophyticus* UTI as was recently shown for *E. coli* (*73*). Additionally, our results show that built environments, including some isolates from wastewater, appear to act as a ‘sink’ where diversity of *S. saprophyticus* is lost. The duration of colonization of these environments is still in question, whether the colonization is transient and if these strains are transmitted back into the source populations may be ascertained using a more thorough sampling approach.

### Limitations and future directions

The results presented here are limited by the biased sampling of *S. saprophyticus* thus far. Isolates from human infections have been heavily sampled while isolates from other important reservoirs of *S. saprophyticus* like animals and the natural environment have been under sampled. In this work we have compensated for differences in sample size wherever possible however future sampling efforts directed at underrepresented sources will increase power for detecting niche-specific adaptations and further clarify the source-sink dynamics at play in *S. saprophyticus* ecology. Combining whole-genome sequencing (WGS) efforts with epidemiological surveys will help elucidate the transmission network of *S. saprophyticus* and inform strategies for the control of human and animal infection by this pathogen.

## Methods

### Isolation & growth of S. saprophyticus

All incubation steps in the isolation protocol are at 37°C with 5% CO_2_ supplementation for 24 hours unless otherwise stated. Wastewater samples taken from the aeration basin were inoculated into Tryptone NN broth (Tryptone [Gibco] with 2 ug/mL novobiocin and 300 ug/mL nalidixic acid) and grown before being struck onto Mannitol Salt Agar (MSA; Neogen cat. NCM0078A) plates. Colonies positive for mannitol fermentation (yellow halo on MSA plates) were gram stained to include only Gram-positive cocci which were then seeded into 3mL LB (Thermo Scientific; cat. 12780052) cultures. Liquid cultures were streaked onto CHROMagar™ Orientation plates (DRG International, cat. RT412) and small, pink, opaque colonies were selected for MALDI-TOF identification performed at the Wisconsin Veterinary Diagnostics Laboratory. Two *S. saprophyticus* isolates from wastewater were identified. Lactose metabolism was assessed by growth on MacConkey agar (Thermo Scientific; cat. CM0007B).

### DNA extraction and sequencing

Isolates from wastewater and isolates provided from collaborators (8 animal/food, 6 environment, 1 human skin, 137 UTI) were grown in tryptic soy broth (TSB; Thermo Scientific; cat. CM0129B) for 24 – 48 hours at 37°C to an OD600 of ∼1. Genomic DNA was extracted using the Qiagen DNeasy kit (cat. 12224-50) and sent to either the University of Wisconsin Madison Biotechnology Center or SeqCoast for library preparation and paired end 150bp sequencing. Raw sequencing data has been deposited to the NCBI SRA under the project accession PRJNA928770.

### De novo genome assembly and annotation

Raw sequencing data were quality-checked and trimmed using FastQC v0.11.8 (*74*) and Trimmomatic v0.39 (*75*), respectively. Potential contamination was identified using Kraken2 (*76*) and samples with significant (>20%) contamination were discarded. Samples with minimal contamination (10-20%) were filtered using KrakenTools (*77*) script “extract_kraken_reads.py” to include only reads originating from *S. saprophyticus.* Contigs were assembled using SPAdes v3.13.1 (*78*). Assemblies were checked for quality using Quast v5.0.2 (*79*) filtering out contigs shorter than 500 bp or with coverage lower than 5x, as well as confirming all assemblies had an *N*_50_ > 50,000 bp. Assemblies were annotated using Prokka v1.14.0 (*80*). Metabolic pathways and enzymes were annotated in our de novo assemblies using Metabolic v4.0 (*26*). Default parameters were used for all programs unless otherwise noted.

### Genome collection

All *S. saprophyticus* WGS entries into the NCBI SRA database (accessed December 12, 2022) with sufficient metadata to determine isolation source were downloaded and assembled as described above. Additionally, any entries into the NCBI Assembly database for which raw data were not available were downloaded. After quality filtering we had total sample of 780 genomes, from the following sources: 538 genomes assembled from SRA data, 154 newly sequenced isolates, 87 assemblies from NCBI and 1 ancient DNA assembly (*11*). Genomes were grouped into one of five isolation sources: animal (mostly farm animals), built environment (human built and occupied spaces), food (food products and food production environments), human (infection and natural colonization) and natural environments (mostly soil and water). This resulted in the following sample: 32 animal, 67 built environment, 105 food, 559 human and 17 natural environment isolates.

### Reference-guided genome assembly

In addition to de novo assembly, we wanted to look at variation within intergenic regions, so we assembled whole genomes against a reference sequence using an in-house pipeline (github.com/myoungblom/RGAPepPipe_MAY). Briefly, raw data was quality checked and trimmed as described above in “De novo assembly and annotation”. Reads were mapped to the *S. saprophyticus* ATCC 15305 reference genome (GCA_000010125.1) using BWA-MEM v0.7.17 (*81*). Samtools v1.17 view and sort (*82*) were used to process SAM and BAM files. Picard v2.26.4 (<u>github.com/broadinstitute/picard</u>) was used to remove duplicates and add read information and Pilon v1.24 (*83*) was used for variant calling. Finally, assembly quality was assessed using Qualimap v2.2.1 BamQC (*84*). For assemblies downloaded from NCBI that did not have raw data, Mummer v4.0.0 (*85*) was used to align the assemblies to the reference genome using a custom script (github.com/myoungblom/sapro_genomics). Repetitive regions were identified in the reference genome using Mummer v4.0.0 (*85*) and these regions were masked in the resulting reference guided alignment.

### Pangenome analyses

Separate pangenome analyses were performed using Roary v3.12.0 (*86*) on the following groups: total sample (n=780), Clade 1 (n=646) and Clade 2 (n=134). For all pangenome analyses the minimum blastp threshold for ortholog clustering was set to 85% (-i 85), paralogs were not split (-s) and a core genome alignment was made using Prank (-e). Rarefaction and accumulation curves were created using modified versions of published scripts (*87*). Briefly, a gene presence-absence matrix was subsampled 100 times without replacement to the desired total number of genomes and the median value for the number of core and pan genes was plotted for each additional genome added to the sample. Scripts for rarefaction and accumulation plots are available here: github.com/myoungblom/sapro_genomics.

### Phylogenetic trees

The core genome phylogeny was inferred using the core genome alignment output by Roary (see “Pangenome analyses”) using RAxML v8.2.3 (*88*) using the general time reversible (GTR) model of nucleotide substitution and the CAT approximation of rate heterogeneity with non-parametric bootstrapping using the ‘autoMR’ convergence criteria. Recombinant regions were removed from the phylogeny using ClonalFrameML (*18*) as described below.

### Horizontal gene transfer

Recombinant fragments were inferred in the core genome using ClonalFrameML (*18*). The core genome alignment was converted into an extended multi-fasta (XMFA) file using a custom script (github.com/myoungblom/sapro_genomics) and run with the core genome phylogeny inferred using RAxML (see above) as the input tree, using 100 simulations (-emsim 100). The output was used to calculate r/m values (https://github.com/xavierdidelot/ClonalFrameML/issues/92), plot recombinant fragments and the recombination adjusted phylogeny was used for all figures. Recombination analyses were performed identically for the full sample and for the two clades separately. Inter-clade recombination events within the core genome alignment were predicted using FastGear (*23*) with default parameters. Recombinant fragments predicted by ClonalFramML and FastGear were plotted in R using custom scripts (github.com/myoungblom/sapro_genomics).

### RMS search

All RMS gene nucleotide sequences were downloaded from REBASE (*25*) (accessed March 17, 2023) and searched against our de novo assemblies using blastn with filters to include only matches with >80% sequence identity and >80% of RMS gene length. For overlapping alignments, the result with the highest bitscore was chosen.

### Plasmid identification

A database of plasmid sequences from PLSDB (*20*) (accessed February 2, 2023) was downloaded and searched against our de novo assemblies using blastn with filters to include only matches with >80% sequence identity and >80% plasmid length. All overlapping alignments were kept for downstream analysis. Putative plasmid sequences were pulled from the de novo assemblies and pairwise mash distances were calculated with Mash v2.2 (*89*). To visualize the sequence relatedness of plasmids in our sample we performed a multi-dimensional scaling (MDS) analysis of mash distances in R using ‘cmdscale’. Plasmid sequences were grouped into eight sequence types based on a plot of the MDS results.

### Diversity & selection statistics

ANI was calculated between all possible alignment pairs using fastANI (*90*) with a masked whole-genome alignment (see “Reference-guided genome assembly”). Pairwise core genome and accessory gene nucleotide diversity (pi) was calculated using EggLib v3.0.0 (*91*) with custom scripts. dN/dS was calculated across core gene alignments using the yn00 implementation (*92*) in PAML (*93*). dI/dS was calculated as previously described (*94*). Briefly, an alignment of core intergenic regions was made using Piggy (*95*) with all the same flags as were used in the pangenome calculation with Roary (see “Pangenome analyses”). dI was calculated by dividing the number of SNPs in the intergenic alignment by the length of the alignment. The dS values calculated from the core genome alignment were used to calculate both dN/dS and dI/dS. Population genetics statistics (Tajima’s D and nucleotide diversity) were calculated using EggLib v3.0.0 (*91*). Scripts for PAML analysis, sliding window and diversity statistics available here: github.com/myoungblom/sapro_genomics.

### Whole genome variant GWAS

pySEER v1.3.11 (*34*) was used to identify variants in the *S. saprophyticus* whole-genome and core genome alignments that are associated with isolation source/niche. Briefly, population structure was accounted for using phylogenetic distances extracted from the core genome phylogeny using the “phylogeny_distance.py” script included with pySEER. Then for each phenotype (isolation source), pySEER was run using a linear mixed model (LMM) using the phylogenetic distances described above, a phenotype file with the isolation sources of all isolates and a VCF file of SNPs from the whole genome alignment made using SNP-sites v2.4.1 (*96*). The mixed model was chosen out of all models implemented in pySEER because it is a top performing model among microbial GWAS tools (*97*) and is computationally efficient for large samples. The “—output-patterns” flag was used to get the number of unique variant patterns, which when used with the “count_patterns.py” script included with pySEER, outputs a significance threshold using a Bonferroni correction. This significance threshold was used to determine the significance of all pySEER output in addition to removing all results with a “bad-chisq” note, which indicates a failed chi-squared test. Mutation consequences (synonymous, non-synonymous, etc) of the significant pySEER results were annotated using SnpEff (*98*) with a custom database produced using the *S. saprophyticus* reference genome and annotation files (GCA_000010125.1). Scripts for pySEER analyses are available at github.com/myoungblom/sapro_genomics.

## Acknowledgements

We would like to thank the following people for providing samples and/or isolates for this project: Kathleen Glass & Kristin Schill (Food Research Institute), Nicole Aulik (Wisconsin Veterinary Diagnostic Laboratory), Kalan Lab (Dept. of Medical Microbiology and Immunology), McMahon Lab (Dept. of Bacteriology), Derrick Chen (UW Hospital – Clinical Microbiology), Jon Bethke (Duke University), and Marylin Roberts (University of Washington). We would also like to thank members of the Pepperell Lab for their work gathering isolates and performing DNA extractions: Lindsey Bohr, Seanna Curran, Aidan MacKnight, Holly Murray, and Tracy Smith. We also thank the University of Wisconsin-Madison Biotechnology Center and SeqCoast for sequencing services.

## Conflicts of interest

The authors declare no conflicts of interest.

## Funding

MAY was funded by National Science Foundation Graduate Research Fellowship Program under grant No. DGE-1747503. This research was also supported by the National Institutes of Health NIAID R01AI113287 to CSP.

## Supplemental Figures & Tables

**Table S1:**
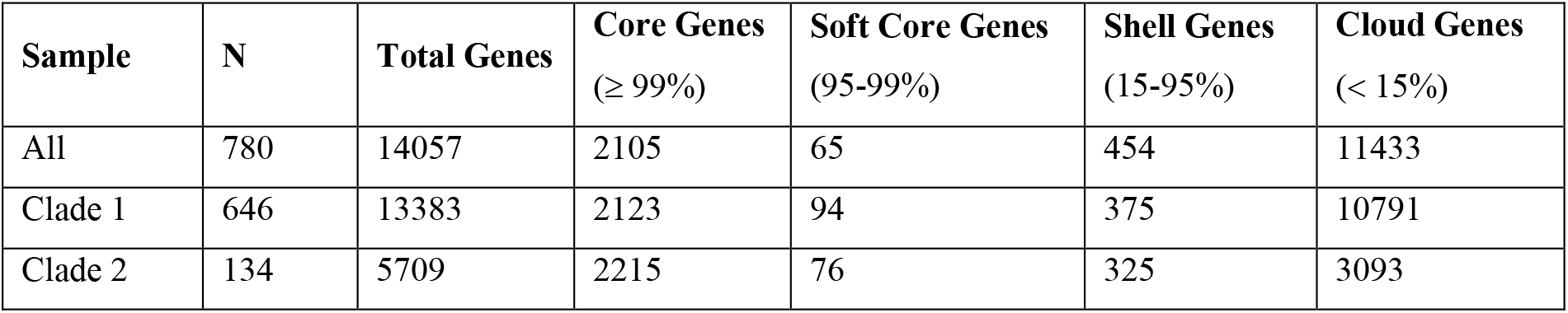
*S. saprophyticus* pangenome dominated by rare accessory gene content. Output of pangenome analyses of full sample, Clade 1 and Clade 2.

| Sample | N | Total Genes | Core Genes<br>(≥ 99%) | Soft Core Genes<br>(95-99%) | Shell Genes<br>(15-95%) | Cloud Genes<br>(< 15%) |
| --- | --- | --- | --- | --- | --- | --- |
| All | 780 | 14057 | 2105 | 65 | 454 | 11433 |
| Clade 1 | 646 | 13383 | 2123 | 94 | 375 | 10791 |
| Clade 2 | 134 | 5709 | 2215 | 76 | 325 | 3093 |

**Figure S1:**
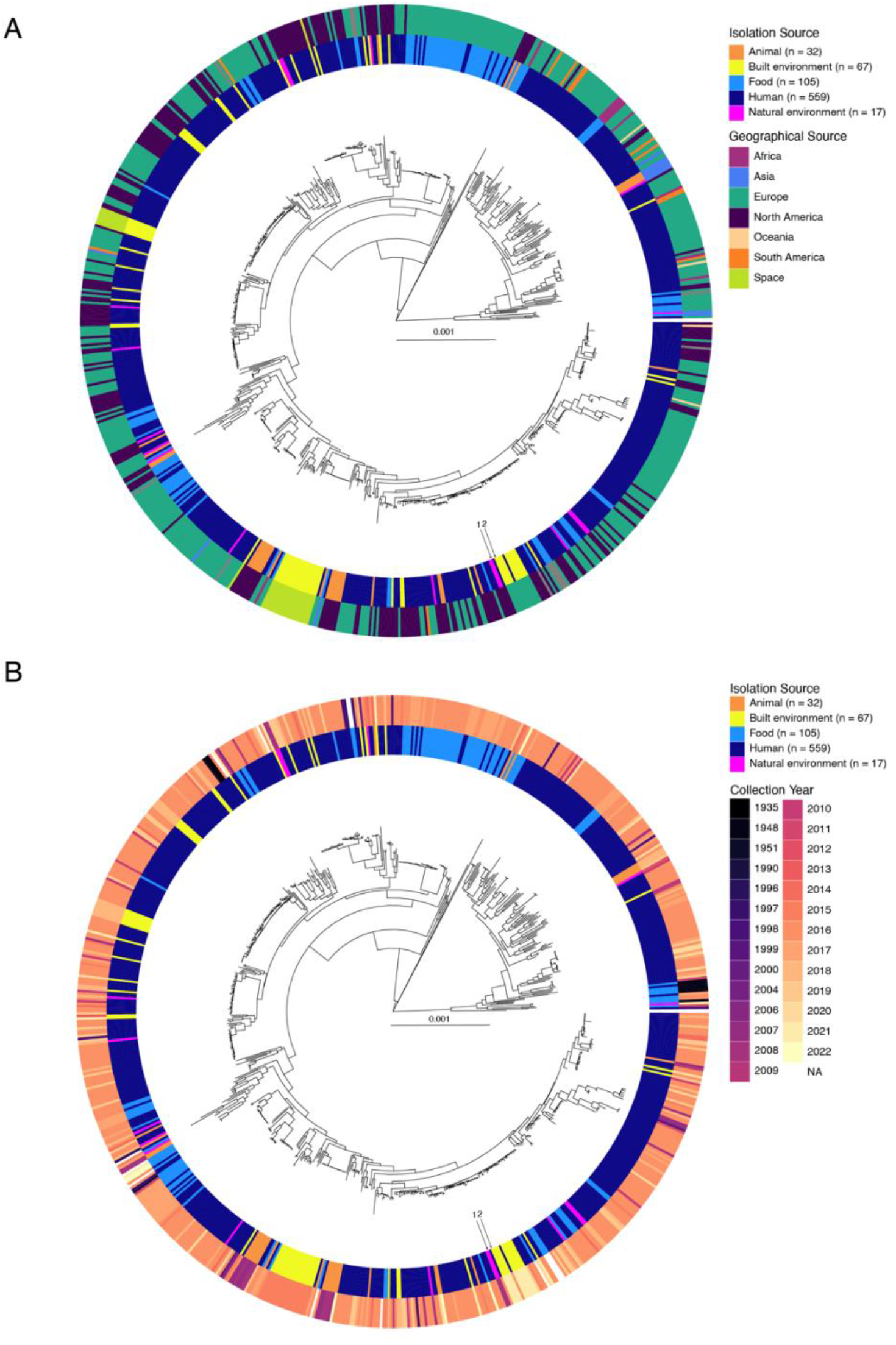
Lack of geographic or temporal structure on the *S. saprophyticus* phylogeny. A) Core genome phylogeny is plotted with isolation source (inner ring) and geographic source (outer ring). B) Core genome phylogeny is plotted with isolation source (inner ring) and isolation year (outer ring). An example of two closely related isolates that illustrate the lack of geographic or temporal structure are marked on A and B: 1) 20-05 (Washington Pacific Ocean, 2008) and 2) SRR19995418 (Norwegian bathroom, 2021) are separated by only 47 core genome SNPS (5e-5 SNPs per site).

**Figure S2:**
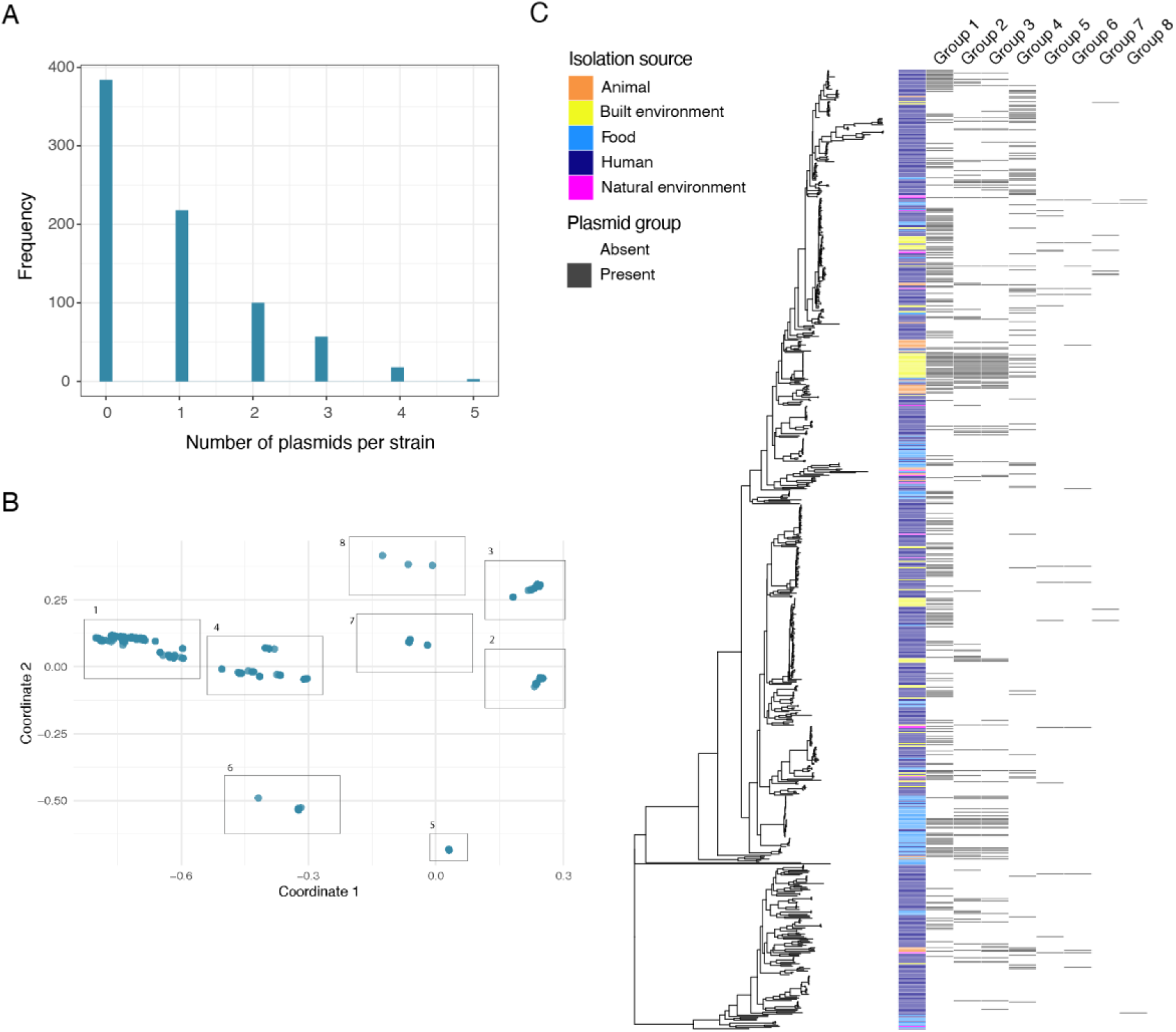
*S. saprophyticus* plasmids are diverse in number, sequence, and phylogenetic distribution. A) Plasmid counts per isolate as determined by the number of unique contigs mapping to plasmids in our plasmid database (see Methods). In about 50% of isolates, nothing resembling any previously sequenced plasmid was identified. The other 50% of isolates had between 1 and 5 plasmids, with only 4/780 strains carrying 5 plasmids. B) Multi-dimensional scaling (MDS) was performed on all putative plasmid sequences and plasmids were grouped into eight sequence groups. C) Plasmid presence/absence broken down by sequence group is plotted alongside the core genome phylogeny illustrating that plasmid sequence groups are generally distributed throughout the phylogeny

**Figure S3:**
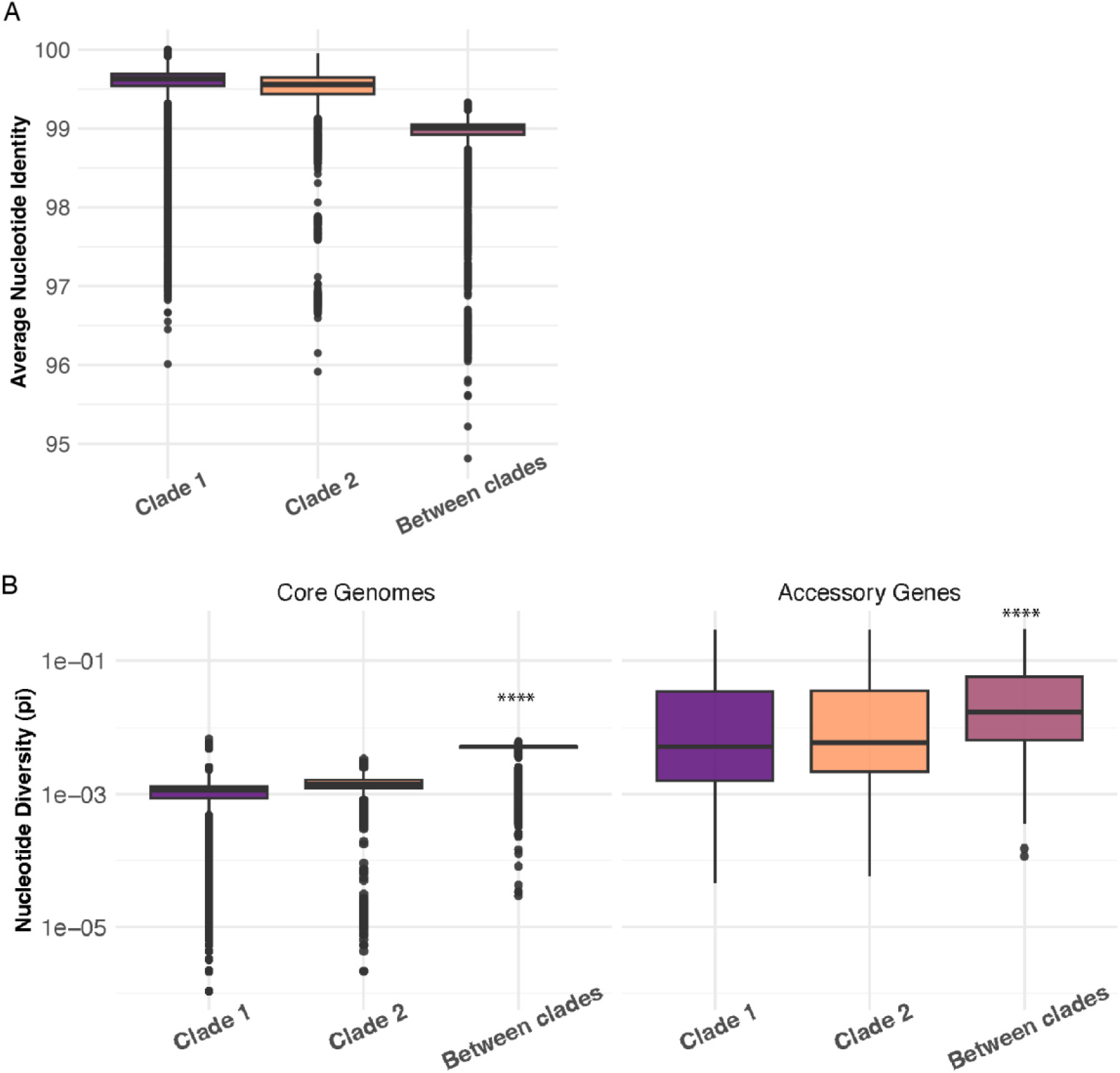
*S. saprophyticus* clades are genetically distinct at the core and accessory genome levels. A) Average nucleotide identity (ANI) calculated from whole-genome alignments. ANI values from isolates of different clades range from 95-99%, which is above the threshold that would distinguish them as different sub-species. B) Nucleotide diversity (pi) calculated for core genomes (left) and accessory genes (right). Higher diversity values of between-clade pairs indicate barriers to horizontal transfer between clades.

**Figure S4:**
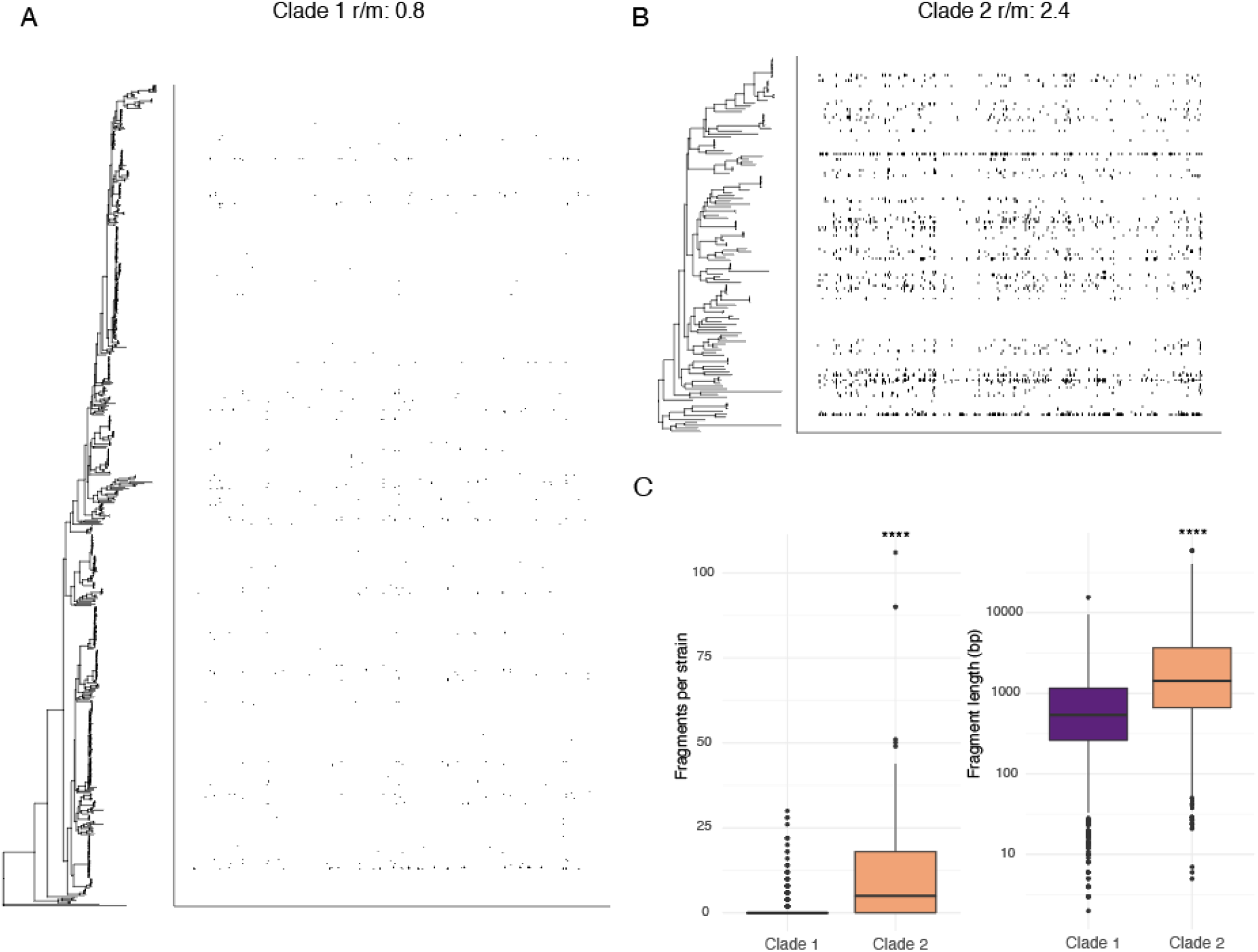
Recombination patterns differ between clades of *S. saprophyticus*. Annotation of recombinant fragments and calculation of r/m values was performed with ClonalFrameML. Plots of recombinant fragments in the core genomes of Clade 1 (A) and Clade 2 (B) show that the core genome of Clade 2 isolates is more affected by recombination. C) Recombinant fragments are significantly more prevalent and significantly longer in Clade 2 isolates (Mann Whitney U test with Bonferroni correction, ****: *p* < 0.0001).

**Figure S5:**
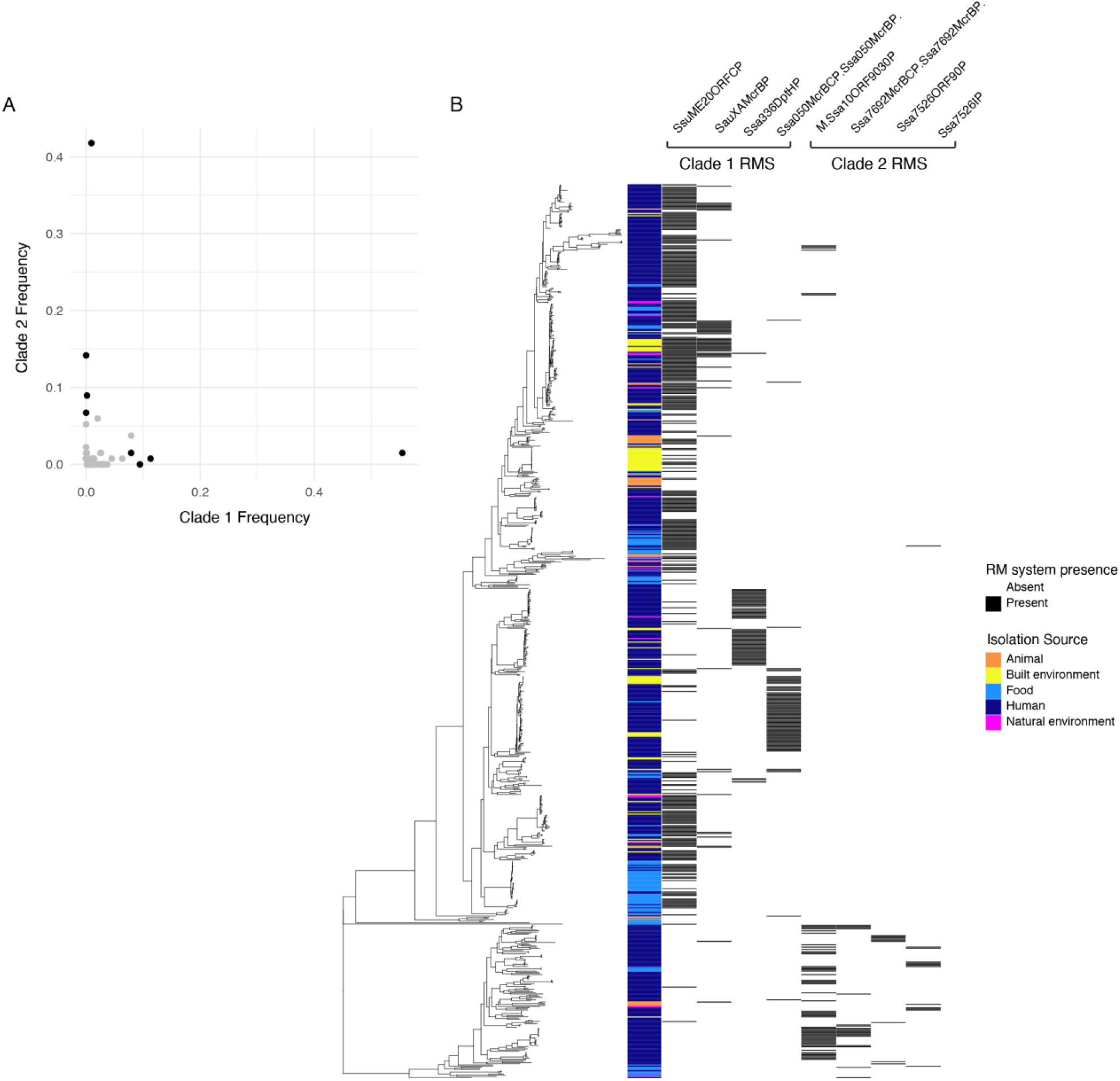
Restriction-modification system genes differ between clades of *S. saprophyticus*. A) Frequency of restriction-modification system (RMS) genes in each clade are plotted. Points representing genes of interest plotted in black. B) Presence/absence of RMS genes of interest plotted alongside the phylogeny showing stark differences in distribution between the two clades.

**Figure S6:**
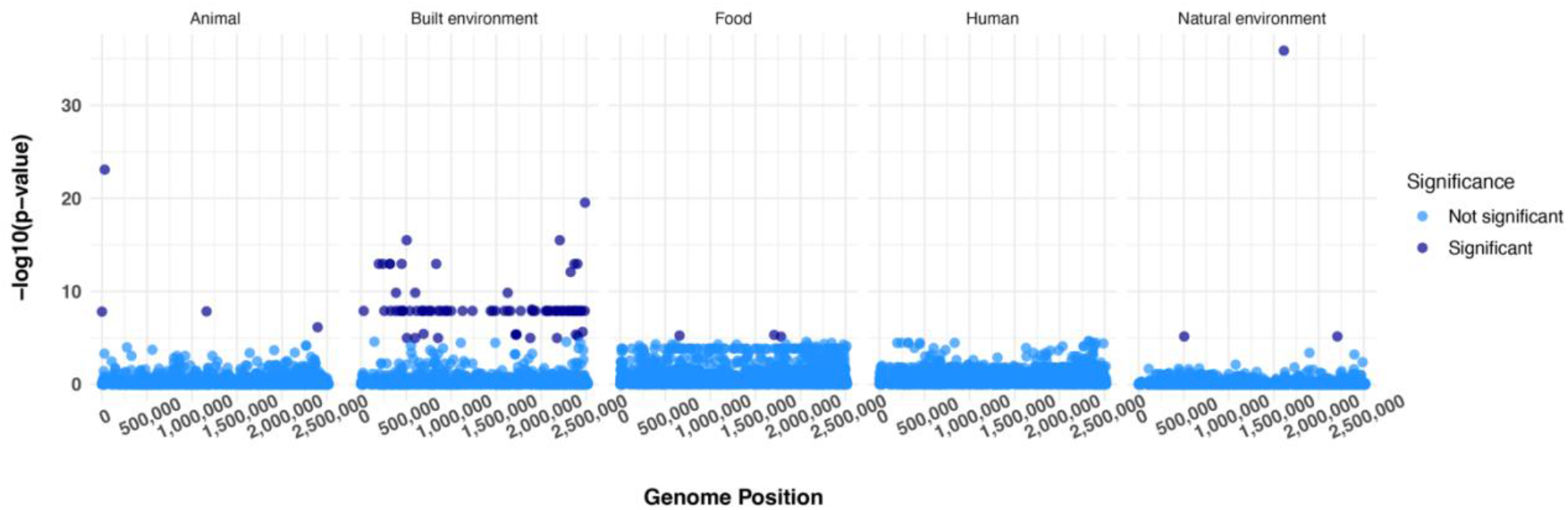
Few core genome variants associated with niche adaptation. GWAS on all SNPs from core genome alignment (n = 6,013) was performed using pySEER with a mixed model. SNPs identified as significantly associated with a particular isolation source are in dark blue, non-significant in light blue. Isolates from built environments have a much higher number of associated variants: 80% of all significant SNPs were associated with built environments.

**Table S2:** Variants from whole-genome alignment significantly associated with isolation source. For intergenic variants, the downstream gene is listed. Gene ID and product refer to the annotations of the reference genome found on NCBI (GCA_000010125.1).

| Position | Association | Mutation Type | Gene ID | COG | Product |
| --- | --- | --- | --- | --- | --- |
| 30938 | Animal | synonymous | SSP0023 | S | 2CRS activity regulator YycH |
| 195852 | Built env. | non-synonymous | SSP0174 | S | nickel pincer cofactor biosynthesis protein LarC |
| 240768 | Built env. | non-synonymous | SSP0212 | - | hypothetical protein |
| 319124 | Built env. | synonymous | SSP0294 | E | aminotransferase |
| 320373 | Built env. | synonymous | SSP0295 | CH | D-2-hydroxyacid dehydrogenase |
| 386784 | Built env. | intergenic | SSP0353 | E | aminotransferase |
| 449952 | Built env. | non-synonymous | SSP0411 | G | gluconokinase |
| 505482 | Built env. | missense | SSP0467 | S | DUF805 domain |
| 600739 | Built env. | synonymous | SSP0564 | S | ABC-type multidrug transport system ATPase |
| 833938 | Built env. | synonymous | SSP0807 | E | alanine racemase |
| 1627287 | Built env. | non-synonymous | SSP1559 | L | primosomal protein |
| 1896024 | Built env. | synonymous | SSP1814 | E | argininosuccinate synthase |
| 2206045 | Built env. | synonymous | SSP2142 | EGP | proline betaine transporter |
| 2326532 | Built env. | synonymous | SSP2252 | S | polysaccharide biosynthesis protein |
| 2368785 | Built env. | non-synonymous | SSP2284 | E | glutamate synthase large subunit |
| 2401511 | Built env. | non-synonymous | SSP2320 | - | hypothetical protein |
| 657433 | Food | intergenic | SSP0618 | P | ABC-type amino acid transport |
| 1708638 | Food | non-synonymous | SSP1643 | S | hypothetical protein |
| 1785389 | Food | non-synonymous | SSP1717 | F | phosphoribosylaminoimidazolecarboxamine formyltransferase |
| 1611892 | Natural env. | synonymous | SSP1545 | E | L-serine dehydratase beta subunit |

**Figure S7:**
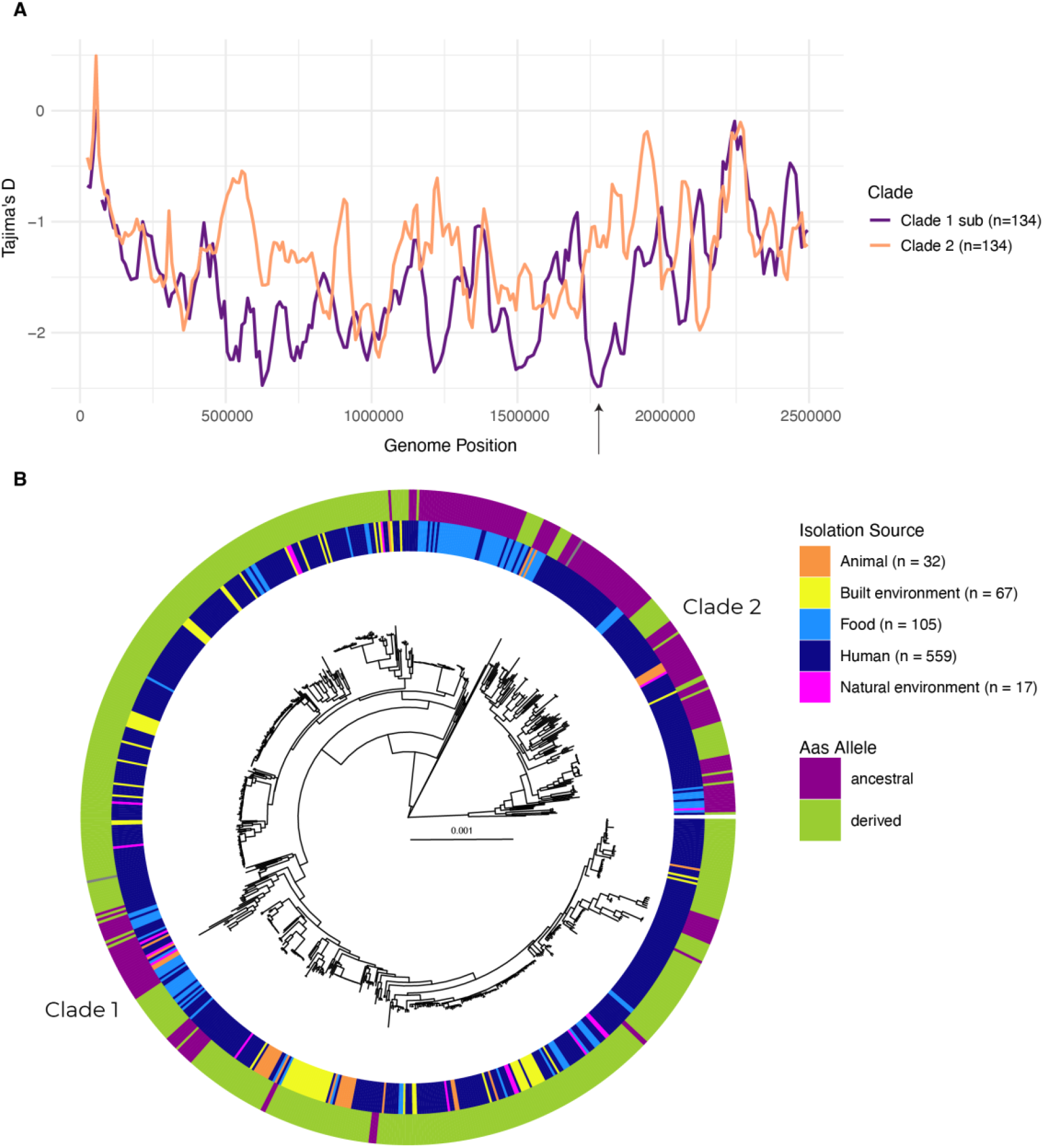
Selective sweep in *aas* associated with Clade 1. A) Tajima’s D calculated in sliding windows (window size: 50,000 bp, step size: 10,000 bp) across whole genome alignments of each clade. Alignments were repeatedly (100x) sub-sampled to the size of the smaller clade (Clade 2, n=134) and the mean Tajima’s D was plotted. Clade 1 has overall lower Tajima’s D (-1.6) than Clade 2 (-1.2). Evidence for a previously identified selective sweep in the bifunctional adhesin-autolysin Aas is replicated in this larger dataset as evidenced by the dip in Tajima’s D in region near 1,775,000 bp (marked by the arrow) of the Clade 1 alignment. B) Alleles of the non-synonymous variant (position 1,811,777) in Aas previously identified as having undergone a selective sweep are plotted on the core genome phylogeny (outer ring) alongside the isolation source (inner ring) of all genomes. Aas alleles are highly structured on the phylogeny with Clade 1 having a higher proportion of derived alleles (82%) than Clade 2 (28%).

**Supplementary Data 1:** Table of genomes used in this study with associated metadata.

**Supplementary Data 2:** Accessory genes significantly associated with each isolation source.

## Notes

### Competing Interest Statement

The authors have declared no competing interest.

